# Toggle-Untoggle: A cell segmentation tool with an interactive user verification interface

**DOI:** 10.1101/2025.05.21.655178

**Authors:** Nina Grishchenko, Margarita Byrsan, Michael F. Olson

## Abstract

Accurate cell segmentation is an essential step in the quantitative analysis of fluorescence microscopy images. Pre-trained deep learning models for automatic cell segmentation, such as Cellpose, offer strong performance across a variety of biological datasets but may still introduce segmentation errors. While training custom models can improve accuracy, it often requires programming expertise and significant time, limiting the accessibility of automatic cell segmentation for many wet lab researchers. To address this gap, we developed “Toggle-Untoggle”, a desktop application that combines automated segmentation using the Cellpose “cyto3” model with a user-friendly graphical interface for intuitive segmentation quality control. Our app allows users to refine results by interactively toggling individual segmented cells on or off without the need to manually edit segmentation masks, and to export morphological data and cell outlines for downstream analysis. Here we demonstrate the utility of “Toggle-Untoggle” in enabling accurate, efficient single-cell analysis on real-world fluorescence microscopy data, with no coding skills required.

## Introduction

Accurate cell segmentation is a necessary step for robust quantitative analysis of fluorescence microscopy images, enabling the identification of individual cells and the extraction of morphological features, such as those related to size, shape, and intensity, which are essential for studying cell behaviors, phenotypic variation, and treatment responses (Li et al., 2013). These morphological features are often used in downstream applications such as clustering and classification, where segmentation quality directly impacts the reliability of the results (Ahmadzadeh et al., 2017; Yao et al., 2019)

Deep learning approaches, particularly convolutional neural networks (CNNs) such as U-Net, have significantly advanced segmentation accuracy in bioimage analysis (Ronneberger et al., 2015). A U-Net style architecture called Cellpose has gained popularity for its generalist pre-trained models that perform well across a range of imaging conditions and cell types (Stringer et al., 2021). The latest Cellpose version introduced the “cyto3” model, which improved segmentation performance while eliminating the need for user-specific model training, making it more accessible to users without deep learning expertise (Stringer & Pachitariu, 2025).

Despite these advantages, pre-trained models are not always well-suited to every dataset. In cases where cell shapes, textures, or imaging conditions markedly differ from the training data, segmentation errors may occur including missing cells, over-segmentation, or poor boundary detection, each of which could compromise the accuracy and robustness of extracted morphological data (Chen & Murphy, 2023). Manual correction is often necessary, which can be slow and difficult when processing large volumes of images.

To address this challenge, we developed “Toggle-Untoggle”, a desktop application (app) with a graphical user interface (GUI) that integrates automated segmentation with a user-friendly correction/verification tool and streamlined extraction of morphological parameters. The app features an interactive interface with a user input panel (**Figure 1A**), built-in guidance, and troubleshooting tips to help minimize segmentation errors. Designed for accessibility, it requires no coding expertise, making it easy for users of all backgrounds to perform high-quality single-cell image analysis. Images in Tiff format can be automatically loaded from a specified directory for batch processing, and segmentation is performed using the Cellpose super-generalist “cyto3” model (Stringer & Pachitariu, 2025). Following segmentation (**Figure 1B**), the app extracts standard morphological features for each detected cell, and enables users to review and refine results by simply toggling individual segmented cells off, removing them from downstream analysis but retaining their position to allow them to be toggled back on again if required. This interactive correction system provides an efficient way to improve segmentation quality without time-consuming manual editing of masks or identification of outliers in downloaded datasets. In addition, the app enables users to export results in formats compatible with widely used analysis platforms (**Figure 1B**). A .csv file containing single-cell morphological parameters, as well as segmentation masks in .roi format, can be saved for further analysis in external tools such as Fiji or QuPath (Bankhead et al., 2017; Schindelin et al., 2012). These features increase the utility of Toggle-Untoggle for users who require more advanced visualization or analysis beyond the app itself.

**Figure 1.**
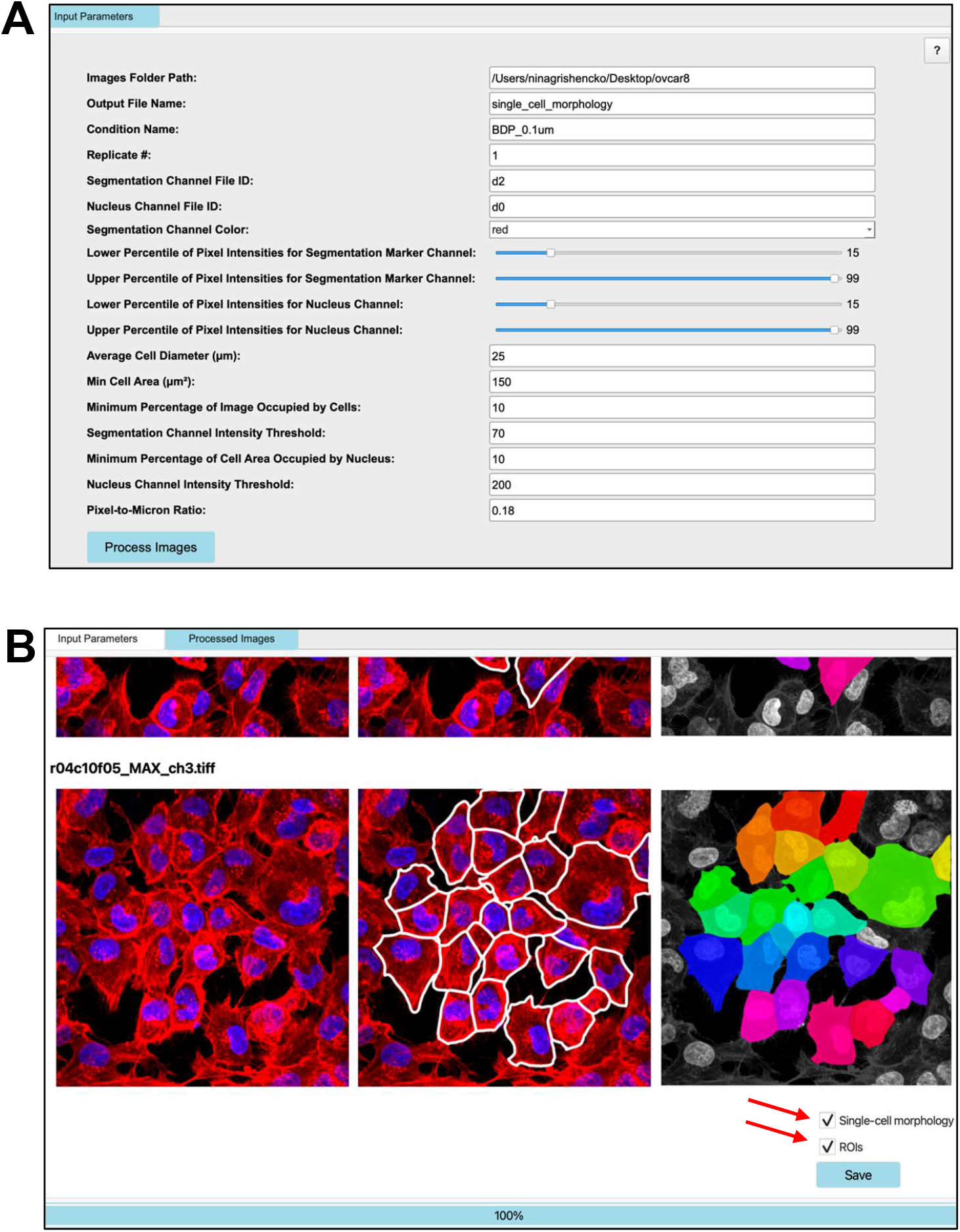
“Toggle-Untoggle” graphical user interface. **A**. An overview of the Input Parameter screen that allows users to upload image datasets and vary conditions for optimal segmentation. **B**. Images of parental OVCAR-8 cells were used as an example segmentation. Fluorescently labelled phalloidin was used to visualize F-actin, which served as the marker for cell segmentation. Red arrows highlight the options located above the ‘Save’ button, allowing users to export a .csv file containing morphological parameters and/or region of interest (ROI) outlines for individual cells of interest.

Here, we present the design and functionality of the application, demonstrate its performance on an image dataset, and highlight its utility in enabling accurate, scalable, and user-friendly single-cell analysis of fluorescence microscopy images of mammalian cells.

## Methods

### Cell culture

Human ovarian endometrioid adenocarcinoma cells (A2780) and a cisplatin-resistant derivative (A2780CisR), obtained from Dr. Robert Rottapel (Princess Margaret Cancer Centre, Toronto Canada), were cultured in RPMI 1640 medium supplemented with 10% fetal bovine serum (FBS), 100 U/mL penicillin, and 100 µg/mL streptomycin. Triple-negative human breast cancer (TNBC) MDA MB 231 cells were cultured in Dulbecco’s Modified Eagle Medium (DMEM) supplemented with 10% FBS, 100 U/mL penicillin, and 100 µg/mL streptomycin. All cells were maintained at 37°C in a humidified incubator with 5% CO_2_.

### Immunofluorescence microscopy

A2780 and A2780CisR cells were seeded at 5,000 cells per well, and MDA MB 231 cells at 12,000 cells per well, in PerkinElmer PhenoPlate 96-well CellCarrier Ultra plates. Cells were cultured for 24 hours at 37°C with 5% CO_2_. After three washes with PBS, cells were fixed with 30 µL/well of 4% paraformaldehyde (PFA) in PBS for 15 minutes at room temperature in the dark. Permeabilization was performed with 50 µL/well of 0.1% Triton X-100 in PBS for 5 minutes, followed by blocking with 50 µL/well of 3% (w/v) bovine serum albumin (BSA) in PBS for 1 hour at room temperature.

A2780 and A2780CisR cells were stained with rhodamine-conjugated phalloidin (1:200, ThermoFisher Scientific) and 4′,6-diamidino-2-phenylindole (DAPI, 1:5000, ThermoFisher Scientific) in 3% BSA for 1 hour to label F-actin and nuclei, respectively. MDA MB 231 cells were stained with rhodamine-conjugated phalloidin (1:200) and DAPI (1:5000) as above, along with an anti-green fluorescent protein (GFP) N-terminal rabbit polyclonal antibody (1:200, Sigma-Aldrich) in 3% BSA for 1 hour. After washing three times with PBS, cells were incubated with Alexa Fluor® 488-AffiniPure Donkey Anti-Rabbit IgG secondary antibody (1:1000, ThermoFisher Scientific) in 3% BSA for 1 hour in the dark. Following three final PBS washes, cells were stored in 50 µL PBS at 4°C in the dark until imaging. High-content imaging was performed using an Opera Phoenix™ Plus high-content imaging system.

### Graphical user interface development workflow

“Toggle-Untoggle” was built with PyQt6. It uses the Cellpose algorithm for cell segmentation and scikit-image to extract morphological parameters of individual objects. The full code and first release of the app are available to view and download on our GitHub page: https://github.com/ninagris/Toggle-Untoggle

### Data analysis and visualization

All statistical comparisons were conducted using Student’s t-test, with a p-value ≤ 0.05 considered statistically significant using GraphPad Prism. Data presentation also used GraphPad Prism. Haralick texture features were extracted and analyzed using QuPath software.

## Results

A custom GUI application was developed with PyQt6 to facilitate automated cell segmentation, morphological feature extraction, and user-guided verification and correction. The app integrates the pre-trained Cellpose “cyto3” model for segmentation, enabling immediate use without the need for custom training. Upon launch, users select a directory containing input images, which are automatically loaded and processed in batch mode (**Figure 1A**). The interface includes adjustable input fields for key parameters, including: the channels to be analyzed; parameters to enhance the appearance of the original false-color images for user reference and assistance with post-segmentation verifications and corrections; estimated cell diameter; minimum cell area to eliminate under-segmented cells and debris; and intensity specifications to filter out empty images and under-segmented objects lacking a nucleus. These are supplemented with on-screen instructions to assist users in optimizing segmentation performance and troubleshooting if necessary (**Figure 1A**).

For each image, Cellpose is applied to generate cell outlines and segmentation masks based on the chosen cytoplasmic marker. These masks are overlaid on the original image and displayed in the GUI for review (**Figure 1B**). After processing all images in the directory, or only a subset using the “stop processing” button, users can review and modify the segmentation results. Exportable .csv files include the morphological parameter data for each cell. Regions of interest (ROIs) for the segmented cells can also be saved for downstream analysis in external tools, if applicable (**Figure 1B**). All output files are saved to the folder containing the input images used for the analysis.

Since pre-trained models may not always yield perfect results, particularly in the presence of cell clusters or artifacts, the application includes the interactive Toggle-Untoggle feature. As shown in the example of MDA MB 231 breast cancer cells that were segmented based on expression of a GFP marker, even after troubleshooting with all the available parameters in the user input panel on the first tab of the app, incorrectly segmented cells may still occur (**Figure 2A**). This can cause users to doubt the accuracy of the segmentation and prompt them to correct mis-segmented objects. As demonstrated in **Figure 2B**, “Toggle-Untoggle” provides a straightforward solution to this commonly encountered problem. The interactive tool built into the GUI allows users to exclude incorrectly segmented cells simply by clicking on them, which increases the opacity of the overlay and ensures that these cells are not included in the exported dataset or saved ROIs. Verifications/corrections can be performed across all images and cells displayed in the GUI, which is then implemented in the exported data. Users have the option to add selected cells back to the segmented cells in each image by clicking on the opaque version, at which point the cell mask will revert to the original settings and the selected cell will be included in the exportable dataset and ROIs. This feature ensures that toggling individual cells to exclude them from analysis is fully reversible (untoggling), should it be determined that their exclusion was in error. Users also have the option to save two .csv files: one containing verified segmented cells, and another containing cells “toggled” or removed by the user. Both files include the morphological parameter data for each cell. The possibility of exporting two separate files containing either the accepted or the removed cells could allow for refinement of automated segmentation algorithms by incorporating training for acceptable and unacceptable cell masks.

**Figure 2.**
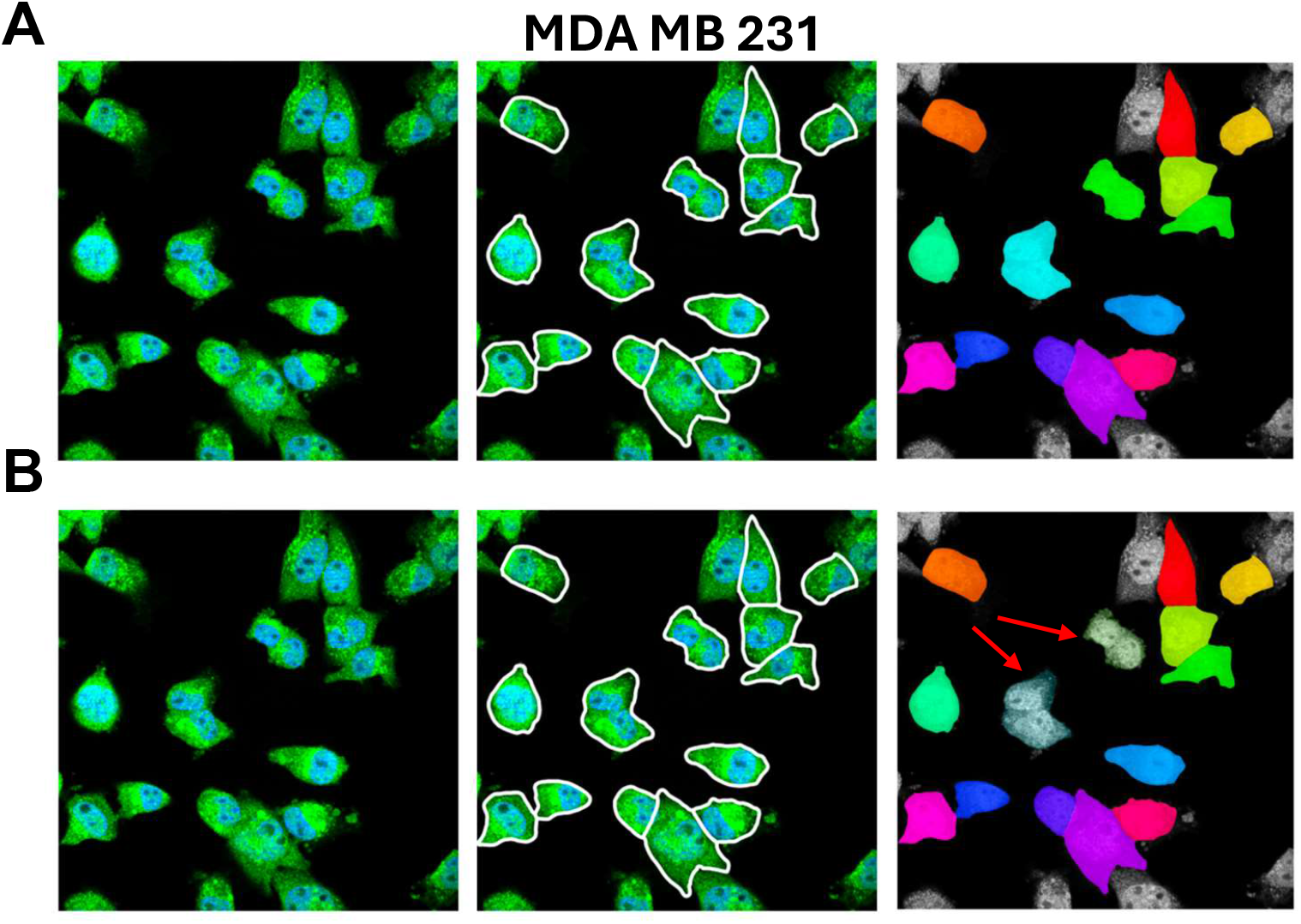
Toggle/Untoggle feature for de-selecting segmented cells. MDA MB 231 cells were segmented based on expression of a GFP marker. **A**. The original segmentation masks can be toggled on or off by the user. **B**. De-selected masks, indicated by the red arrows, are made opaque and excluded from the final dataset.

Following segmentation and optional verification/correction, single-cell morphological parameters are automatically determined using the scikit-image library. Some of the extracted features include area, perimeter, eccentricity, solidity, major and minor axis lengths, and mean fluorescence intensity. These parameters provide quantitative descriptions of cell shape and signal, enabling detailed comparisons between different experimental conditions. Here, we demonstrate how the functionality of “Toggle-Untoggle” was used to compare the morphology of parental and cisplatin-resistant OVCAR8 cells, which appear differ in size from a qualitative perspective. We segmented A2780 (**Figure 3A**) and A2780CisR (**Figure 3B**) cells based on their F-actin structures that were visualized with fluorescent phalloidin. Using the quantitative single-cell morphological data acquired with “Toggle-Untoggle,” we then performed statistical analyses to compare the appearance of parental and cisplatin-resistant A2780 cells. Cell area (**Figure 4A**), mean phalloidin fluorescence intensity (**Figure 4A**), and cell perimeter (**Figure 4C**) were each statistically significantly different between parental A2780 and cisplatin-resistant A2780CisR ovarian cancer cells.

**Figure 3.**
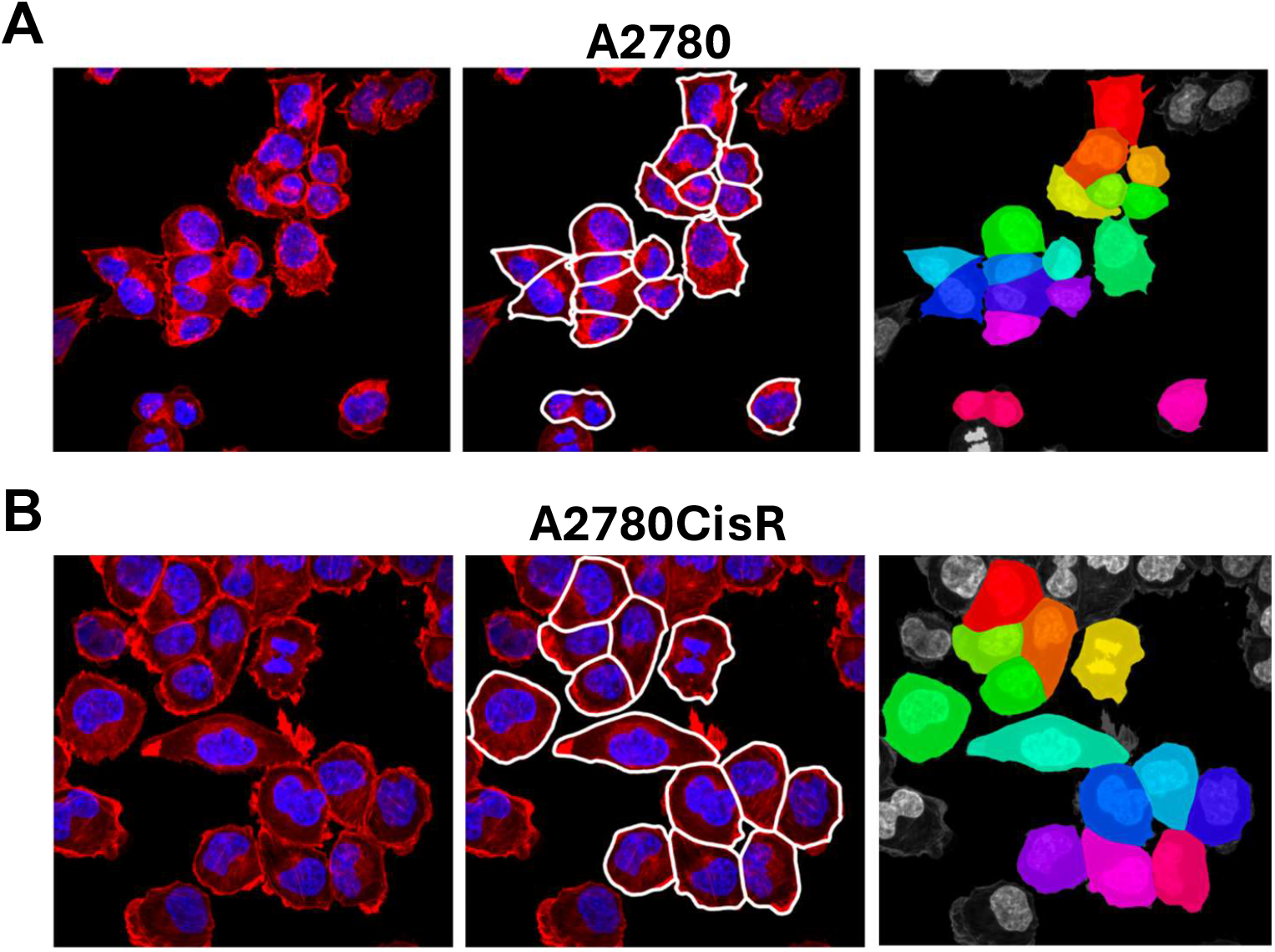
Segmentation of parental and cisplatin-resistant A2780 cell lines. **A**. Parental A2780 and **B**. cisplatin-resistant A2780CisR cell lines were segmented based on phalloidin staining of F-actin structures. Images were analyzed using the “Toggle-Untoggle” GUI to extract single-cell morphological parameters and regions of interest.

**Figure 4.**
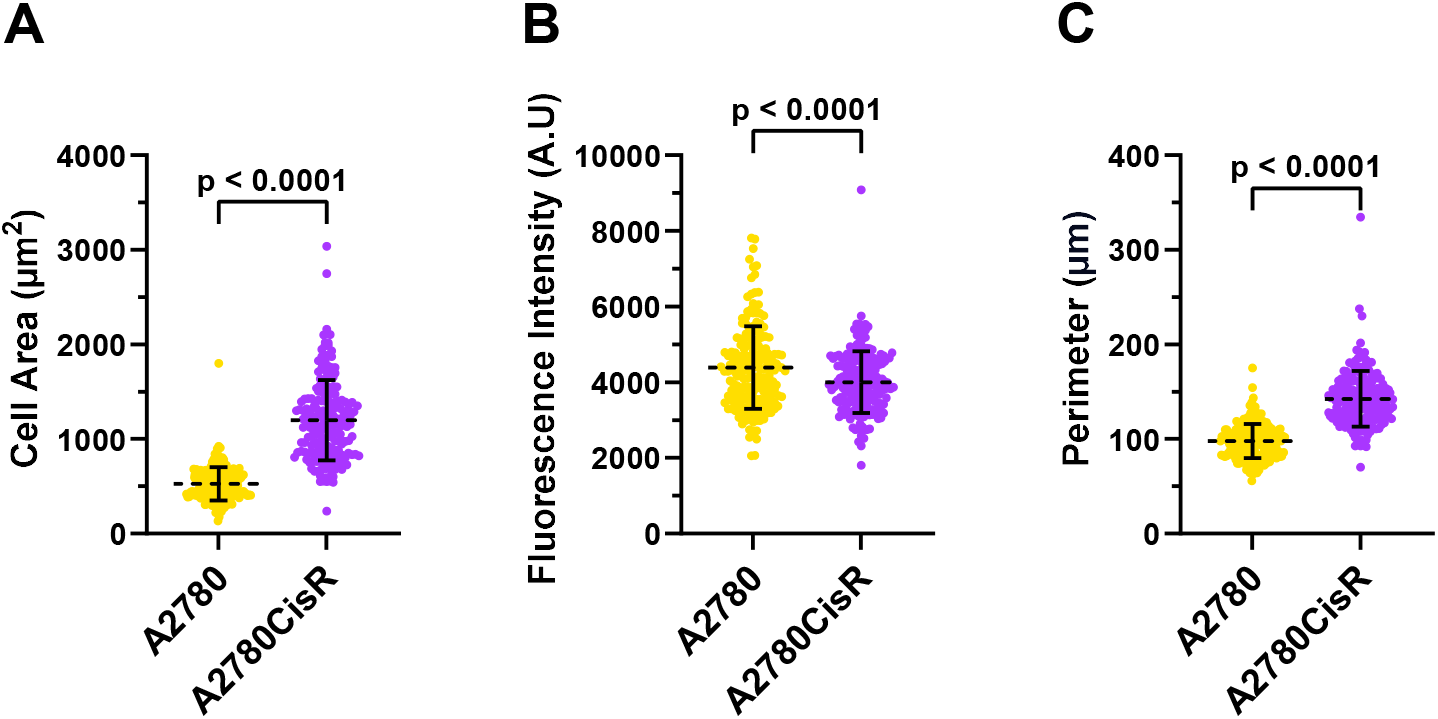
Comparative morphological analysis of parental and cisplatin-resistant A2780 Cells. Cell segmentation and feature extraction for parental A2780 and cisplatin-resistant A2780CisR cells were performed using “Toggle-Untoggle”, enabling quantification of **A**. cell area, **B**. mean fluorescence intensity, and **C**. cell perimeter. Statistical analyses were conducted using Student’s t-test, with a sample size of *n* = 200 cells per group.

To demonstrate the feasibility of downstream analysis using the cell segmentation mask produced by “Toggle-Untoggle,” the ROI masks generated by the app were saved and imported into the third-party platform QuPath (Bankhead et al., 2017) to extract Haralick texture features, enabling an evaluation of differences in actin texture between the parental and cisplatin-resistant A2780 cell populations (**Figure 5**). Haralick features can be used to quantify textural properties in images by capturing patterns of pixel intensity variation, which can reflect underlying biological structures such as the organization of the actin cytoskeleton (Hamilton, 2009; Hui et al., 2023). In this study, we determined that specific Haralick features were statistically significantly different between parental A2780 and cisplatin-resistant A2780CisR ovarian cancer cells, including: contrast, which measures the extent of local variations present in the image (**Figure 5A**); inverse difference moment (IDM), which captures local homogeneity by weighting neighboring pixel pairs with similar intensities more heavily (**Figure 5B**); sum average, representing the average sum of pixel pairs (**Figure 5C**); entropy, reflecting the degree of randomness in pixel intensity (**Figure 5D**); difference entropy, describing the randomness in differences between neighboring pixel intensities (**Figure 5E**); and difference variance, which quantifies the variation of intensity differences (**Figure 5F**) (Haralick et al., 1973; Soh & Tsatsoulis, 1999). Together, these metrics provide complementary information about cytoskeleton and organization, offering insights into structural changes associated with drug resistance.

**Figure 5.**
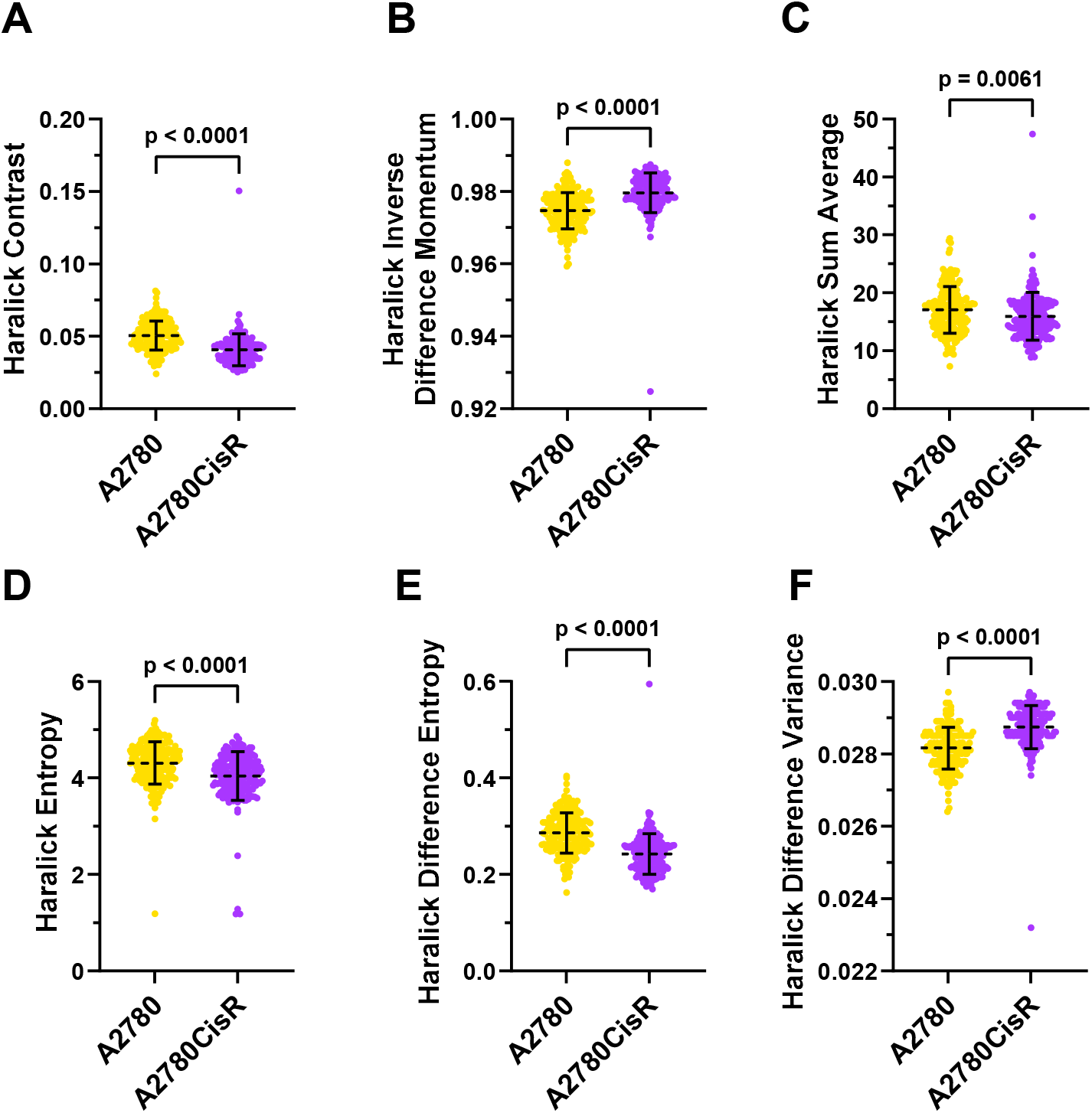
Haralick texture features in parental and cisplatin-resistant A2780 cells. Regions of interest (ROIs) for parental A2780 and cisplatin-resistant A2780CisR cells were exported from “Toggle-Untoggle” and processed using QuPath to produce Haralick texture features. Selected features displayed include **A**. Haralick contrast, **B**. inverse difference moment, **C**. sum average, **D**. entropy, **E**. difference entropy, and **F**. difference variance. Statistical comparisons were performed using Student’s *t*-test, with sample sizes of *n* = 198 (A2780) and *n* = 200 (A2780CisR) cells.

## Discussion

Accurate segmentation is a key requirement for reliable single-cell microscopy image analysis (Li et al., 2013). While generalist models such as Cellpose (Stringer et al., 2021) have been developed to work across a variety of cell types without the need for users to train their own models, the results may still be suboptimal and require manual correction. Here, we introduce “Toggle-Untoggle,” a tool designed to assist non-specialist researchers in analyzing fluorescence microscopy images at the single-cell level.

In comparison to existing tools, our approach offers unique advantages tailored to wet-lab workflows. Although Cellpose itself includes a GUI, it is mainly intended for parameter optimization and does not support bulk image analysis (Stringer & Pachitariu, 2025). While comprehensive platforms such as Fiji (Schindelin et al., 2012) and QuPath (Bankhead et al., 2017) provide plugins compatible with Cellpose for batch processing, they often require at least some degree of coding proficiency and do not offer interactive options for correcting segmentation masks, a key feature of “Toggle-Untoggle”.

Nevertheless, our method has certain limitations. While manual correction of segmentation masks can enhance accuracy, it may introduce some level of subjective bias if not systematically reviewed. In addition, due to the time required for user intervention and the current partial reliance on Mac CPUs, “Toggle-Untoggle” is best suited to low-throughput analyses rather than high-content, large-scale screening applications. At this stage, the tool has been developed for MacOS, with plans underway to expand compatibility to other operating systems in future updates.

## Acknowledgements

This research was supported by funding to M.F.O from the Canadian Institutes of Health Research (PJT-169125), Natural Sciences and Engineering Research Council of Canada (RGPIN-2020-05388), and Canada Research Chairs Program (950-231665). A2780 and A2780CisR cells were a kind gift from Robert Rottapel (Princess Margaret Cancer Centre, Toronto Canada).

## Author contributions

Study conception and design N.G. and M.F.O. Program coding N.G. Cell biology and imaging N.G. and R.B. Data analysis and visualization N.G and M.F.O. Manuscript writing, review and editing, N.G. and M.F.O.

